# Allelic gene conversion softens selective sweeps

**DOI:** 10.1101/2023.12.05.570141

**Authors:** Daniel R. Schrider

**Affiliations:** Department of Genetics, University of North Carolina, Chapel Hill, NC 27599

## Abstract

The prominence of positive selection, in which beneficial mutations are favored by natural selection and rapidly increase in frequency, is a subject of intense debate. Positive selection can result in selective sweeps, in which the haplotype(s) bearing the adaptive allele “sweep” through the population, thereby removing much of the genetic diversity from the region surrounding the target of selection. Two models of selective sweeps have been proposed: classical sweeps, or “hard sweeps”, in which a single copy of the adaptive allele sweeps to fixation, and “soft sweeps”, in which multiple distinct copies of the adaptive allele leave descendants after the sweep. Soft sweeps can be the outcome of recurrent mutation to the adaptive allele, or the presence of standing genetic variation consisting of multiple copies of the adaptive allele prior to the onset of selection. Importantly, soft sweeps will be common when populations can rapidly adapt to novel selective pressures, either because of a high mutation rate or because adaptive alleles are already present. The prevalence of soft sweeps is especially controversial, and it has been noted that selection on standing variation or recurrent mutations may not always produce soft sweeps. Here, we show that the inverse is true: selection on single-origin *de novo* mutations may often result in an outcome that is indistinguishable from a soft sweep. This is made possible by allelic gene conversion, which “softens” hard sweeps by copying the adaptive allele onto multiple genetic backgrounds, a process we refer to as a “pseudo-soft” sweep. We carried out a simulation study examining the impact of gene conversion on sweeps from a single *de novo* variant in models of human, *Drosophila*, and *Arabidopsis* populations. The fraction of simulations in which gene conversion had produced multiple haplotypes with the adaptive allele upon fixation was appreciable. Indeed, under realistic demographic histories and gene conversion rates, even if selection always acts on a single-origin mutation, sweeps involving multiple haplotypes are more likely than hard sweeps in large populations, especially when selection is not extremely strong. Thus, even when the mutation rate is low or there is no standing variation, hard sweeps are expected to be the exception rather than the rule in large populations. These results also imply that the presence of signatures of soft sweeps does not necessarily mean that adaptation has been especially rapid or is not mutation limited.

## 1 Introduction

For a century, the field of population genetics has sought to characterize the evolutionary forces shaping patterns of genetic diversity within and between species (Fisher, 1923; Haldane, 1924; Wright, 1931). Perhaps the most-widely studied and debated topic in this area has been the evolutionary impact of natural selection (Kimura *et al*., 1968; King and Jukes, 1969; Langley and Fitch, 1974; Gillespie, 2000). Although it is now clear that natural selection plays a large role in shaping patterns of genetic diversity (Begun and Aquadro, 1992; Hahn, 2008; Nordborg *et al*., 1996; McVicker *et al*., 2009; Corbett-Detig *et al*., 2015; Kern and Hahn, 2018), the prominence of positive selection, which drives adaptation by favoring mutations that confer a fitness benefit, remains more controversial (Jensen *et al*., 2019). Some researchers have posited that positive selection has a negligible effect on genetic variation (Lohmueller *et al*., 2011; Hernandez *et al*., 2011; Pouyet *et al*., 2018), while others have argued that its impact may be substantial in at least some organisms (Begun *et al*., 2007; Sattath *et al*., 2011; Langley *et al*., 2012; Enard *et al*., 2014; Booker and Keightley, 2018). It is likely that the support for either claim may differ depending on the study system in question. For example, when considering the rates of molecular evolution in protein-coding sequences, the prevalence of positive selection appears to vary markedly across species (Kern and Hahn, 2018), although this may be due at least in part to differences in the rate of deleterious rather than adaptive substitutions (Galtier, 2016). One manner to investigate the role of positive selection is to search for its population genetic signatures that are left as “footprints” that persist for some time after selection as acted. For example, a selective sweep, in which a beneficial mutation rapidly spreads through a population, will result in a localized reduction in diversity around the selected site (Maynard Smith and Haigh, 1974; Kaplan *et al*., 1989) along with other characteristic skews in patterns of diversity (see below). In recent decades, a variety of statistical tests have been devised to detect the signatures of completed (Fay and Wu, 2000; Kim and Stephan, 2002; Kim and Nielsen, 2004; Nielsen *et al*., 2005; Chen *et al*., 2010) or ongoing (Hudson *et al*., 1994; Sabeti *et al*., 2002; Voight *et al*., 2006; Ferrer-Admetlla *et al*., 2014; Akbari *et al*., 2018) selective sweeps. These methods have enabled researchers to uncover genomic loci responsible recent adaptations in a number of organisms (e.g. Sabeti *et al*. (2002); Voight *et al*. (2006); Qanbari *et al*. (2014); Garud *et al*. (2015); Love *et al*. (2023); Kang *et al*. (2021); Wang *et al*. (2021).

In a classic selective sweep (Figure 1A), now often referred to as a hard sweep, the selected mutation arises in a single individual and is immediately beneficial; thus selection begins at frequency 1*/ψN* (where *N* is the number of individuals in the population and *ψ* is the ploidy level). Because individuals harboring the beneficial allele leave more ancestors on average than those lacking it, the allele quickly rises in frequency until it eventually reaches fixation (population frequency *p* = 1.0); upon fixation, at the selected site, every individual traces their ancestry back to a single individual at a very recent time in the past (i.e. no further back than the individual in which the mutation first occurred). This essentially wipes out genetic variation in the immediate genomic vicinity of the selected site, and at linked loci produces an excess of both high- and low-frequency neutral alleles (Braverman *et al*., 1995; Fay and Wu, 2000; Kim and Stephan, 2002) as well as an excess of linkage disequilibrium (Kelly, 1997) on either side of the selected site (Kim and Nielsen, 2004). At increasing genetic distances from the selected site, these signatures dissipate and diversity recovers to background levels.

**Figure 1:**
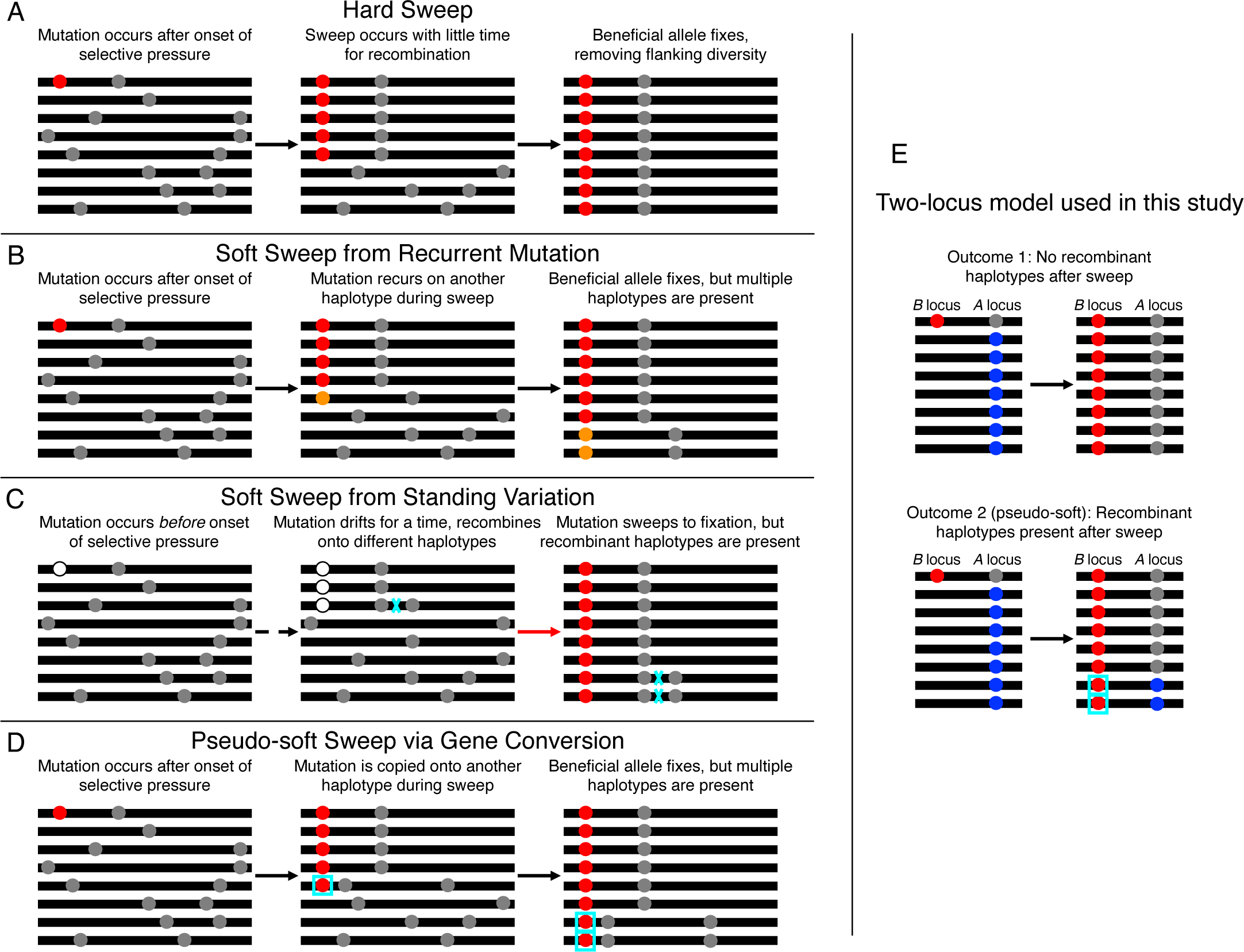
Illustrations of hard and soft sweeps, and the two-locus model used in this study. A) Illustration of a hard sweep occurring from a single-origin *de novo* mutation. Black bars represent different chromosomes present in the population, with circles representing mutations. The red circle is the beneficial mutation, and the gray circles are neutral mutations. The arrows show the passage of time during the sweep. B) A soft sweep via multiple origins of the beneficial allele. Here the first origin of this allele is shown in red, and a recurrent mutation to this allele is shown in orange. At the end of the sweep, there are multiple haplotypes present. C) A soft sweep on a standing genetic variant. Here, the mutation occurs prior to the onset of selection, and is shown as a white circle during this period. The mutation evolves under drift for a period of time (represented by the dashed arrow), before a change in the selective environment causes it to become beneficial and to begin to sweep (red arrow). The mutation (now shown as a red circle) quickly reaches fixation, but multiple haplotypes are present because of recombination events (cyan “x”) and mutation events (not shown) that occurred during the drift phase. D) A pseudo soft-sweep resulting from a single-origin *de novo* mutation followed by allelic gene conversion. During the sweep, the beneficial allele is copied onto another haplotype by a gene conversion event (cyan box) that in this case does not encompass any flanking polymorphisms. Thus, at the end of the sweep there are multiple independent haplotypes present, similar to a soft sweep via recurrent mutation. E) A diagram of the two-locus simulation model examined in this study to assess the frequency of the pseudo-soft sweep via gene conversion scenario shown in panel D. A beneficial mutation to the *B* allele (red circle) occurs at the *B* locus, and is linked to the gray (or *A*) allele at the *A* locus. Initially, all 2*N −* 1 individuals that do not have the beneficial allele at the first locus have the blue (or *a*) allele at the second locus. No crossovers or mutations except the initial origin of the *B* allele are allowed. However, gene conversion events encompassing the *B* locus are allowed. Gene conversion events at the *A* locus are disallowed. Thus, after fixation, the only way that *aB* haplotypes can be present is if they were created via a gene conversion event that replaced the *b* allele with the *B* allele.

More recently, an alternative model of a selective sweep, called a soft sweep, has emerged (Orr and Betancourt, 2001; Hermisson and Pennings, 2005; Przeworski *et al*., 2005; Pennings and Hermisson, 2006b,a; Karasov *et al*., 2010). A soft sweep contrasts with the hard sweep model in that multiple ancestral individuals harboring the selected allele at the onset of selection leave descendants that are present upon fixation (Hermisson and Pennings, 2017). There are several mechanisms by which positive selection can result in a soft sweep. One is selection on a previously segregating polymorphism (Figure 1B) that was selectively neutral (Hermisson and Pennings, 2005) or slightly deleterious (Orr and Betancourt, 2001) but became beneficial following a change in the selective environment. Another mode is selection on an allele that arises multiple times independently in the same population (Hermisson and Pennings, 2005; Pennings and Hermisson, 2006b,a; Figure 1C). A third mode that is somewhat similar to the recurrent mutation mode is when the selected allele enters the population repeatedly via migration from a donor population (Pennings and Hermisson, 2006b). Note that the delineation between hard and soft sweeps is based on the outcome of the sweep rather the process that generated it: if at the time of fixation the advantageous allele has only a single ancestor that was present at the onset of the sweep, then it is considered to be a hard sweep; otherwise, it is a soft sweep regardless of the selective scenario responsible for it.

The signatures left by soft sweeps differ from those of hard sweeps in several key ways. Most notably, soft sweeps cause a less severe reduction in diversity (Innan and Kim, 2004), an increase in linkage disequilibrium and the proportion of high-frequency derived alleles in the direct vicinity of the selected site rather than in flanking regions (Schrider *et al*., 2015; Kern and Schrider, 2018), and often the presence of multiple haplotypes at intermediate-to-high frequencies in contrast to hard sweeps which tend to yield a single dominant haplotype (Pennings and Hermisson, 2006b; Garud *et al*., 2015; Schrider *et al*., 2015; Harris and DeGiorgio, 2020). Recent studies have found evidence that soft sweeps may be relatively common (Garud *et al*., 2015; Feder *et al*., 2016; Schrider and Kern, 2017; Mughal and DeGiorgio, 2019; Xue *et al*., 2021), although these claims have proved contentious (Harris *et al*., 2018; Schrider and Kern, 2018; Feder *et al*., 2021; Garud *et al*., 2021; Johri *et al*., 2022).

There are several conditions under which one might expect soft selective sweeps to be common. Soft sweeps from recurrent mutation are expected be prevalent when, at the time of selection, a population is very large and/or has a high rate of mutation to adaptive alleles (Hermisson and Pennings, 2005; Karasov *et al*., 2010). Soft sweeps on standing variation will predominate in highly diverse populations, because at the time of the shift in the selective environment there will already be a large amount of standing variation for selection to act on. Such diversity may be a result of a large historical effective population size, a large rate of mutation to adaptive alleles (either due to a large spontaneous mutation rate or a large mutational target size for the selected trait), or both (Hermisson and Pennings, 2017). Either of these types of soft sweeps would be a sign of a rapidly evolving population, where adaptive mutations occur very quickly or indeed are already present at the onset of a new selective pressure. However, there is another mechanism that produces selective sweeps that appear to be soft, and could potentially occur in populations that do not meet the above conditions: allelic gene conversion. Gene conversion, which causes a small tract of DNA from a donor chromosome to replace the homologous stretch on the other chromosome, is the outcome of a sizeable subset of recombination events (Gay *et al*., 2007). If the gene conversion tract happens to encompass a sweeping allele, then that allele may jump onto a different background. Such an event would not technically result in a soft sweep, as upon fixation the selected allele itself has only a single ancestor that was present at the time of selection (Figure 1D). However, the gene conversion tract is typically quite small (*<*1 kb; see Hilliker *et al*., 1994; Wijnker *et al*., 2013; Comeron *et al*., 2012; Williams *et al*., 2015), so the outcome is very similar to a soft sweep via recurrent mutation given that in most cases when a gene conversion tract spans the selected allele, very few linked polymorphisms will be affected, if any. Previously, Jones and Wakeley (2008) incorporated gene conversion into McVean’s (2007) three-locus model for examining LD spanning a selected site, and found that gene conversion may often cause sweeping alleles to jump to new genetic backgrounds, resulting in substantial linkage disequilibrium between the two flanks of the sweep, contrary to expectations under a hard sweep model without gene conversion (Kim and Nielsen, 2004; McVean, 2007). Further, Schrider et al. previously showed that the patterns of diversity produced by hard sweeps with gene conversion may be similar to those generated under a model of selection on standing variation (Schrider *et al*., 2015). Given that gene conversion events can occur at a high rate (Hilliker *et al*., 1991; Comeron *et al*., 2012; Wijnker *et al*., 2013; Williams *et al*., 2015; Miller *et al*., 2016), these studies raise the possibility that gene conversion may indeed would-be cause hard sweeps on single-origin *de novo* mutations to instead generate signatures essentially indistinguishable from soft sweeps, a phenomenon we refer to as pseudo-soft sweeps.

Here, we examine the question of whether pseudo-soft sweeps generated by gene conversion would be expected to occur frequently under realistic demographic scenarios with species-appropriate gene conversion rates. We investigate this phenomenon using two-locus forward simulations under a variety of selective and demographic scenarios (with the latter modeled using estimates from humans, *Drosophila melanogaster*, and *Arabidopsis thaliana*. We describe the parameter combinations that will cause sweeps on single-origin mutations to produce pseudo-soft sweeps, assess the impact of gene conversion on the amount of haplotypic diversity present after the sweep, and discuss the implications of our findings for natural populations.

## 2 Methods

### 2.1 Simulation model and SLiM implementation

We sought to model the process of selective sweeps from *de novo* mutations experiencing gene conversion in 3 different species: humans, *Drosophila melanogaster*, and *Arabidopsis thaliana*. Figure 1D illustrates our model of a selective sweep followed by gene conversion transferring the adaptive allele onto additional genetic backgrounds. To investigate the prevalence of these sweeps, we performed forward-in-time simulations of a model with two loci: one locus experiencing direct positive selection, and one linked locus where, when the beneficial mutation occurs, all individuals with the ancestral allele (*b*) at the selected locus have the *a* allele at the linked neutrally evolving locus, and the individual with the favored allele (*B*) has the *A* allele at the linked locus. Recurrent mutation is disallowed at these two loci, as are crossover events. Gene conversion is allowed at the selected locus only, and this is the only manner in which an *aB* haplotype can arise during the simulation. The *B* locus thus serves as a marker for whether adaptive allele has been shuffled onto multiple genetic backgrounds via gene conversion, and if at the end of the simulation there is at least one *aB* haplotype present in the population (or sample), the sweep is said to be a pseudo-soft sweep.

The relevant parameters of our model include the per-base pair gene conversion initiation rate, the distribution of gene conversion tract lengths, the population size history, and parameters governing the timing and strength of selection. Although it can be quite difficult to estimate gene conversion tract lengths (Miller *et al*., 2012), for our purposes the key quantity is the probability that a given site will undergo gene conversion in a single meiosis. Our model therefore only allows gene conversion at the selected site, and thus we only require a parameter governing the total per-base pair gene conversion rate—this composite parameter is equal to the product of the tract initiation rate and mean tract length. We examined several per-base pair gene conversion rates for each species, each expressed relative to a baseline conversion rate. We set this baseline rate to 500*c*, where *c* is the average crossover rate per base pair estimated for a species. We used the following *c* values which we obtained from `stdpopsim` version 0.2.0 (Adrion *et al*., 2020; Lauterbur *et al*., 2023a): 1.30981 *×* 10*^−^*^8^ for humans (the mean value on chromosome 12 using data from Consortium (2007)), 1.7966*e ×* 10*^−^*^8^ for *D. melanogaster* (the mean on chr3L using data from Comeron *et al*. (2012)), and 8.06452 *×* 10*^−^*^10^ for *A. thaliana*(the genome-wide average rate from Huber *et al*. (2014)). Additional conversion rates examined were all larger than the baseline rate: if we express the per-base pair gene conversion rate as 500 *× g × c*, the values we simulated correspond to *g* = 1 (baseline), *g* = 5, *g* = 10, and *g* = 50. In other words, if we assume an average gene conversion tract length of 500, our gene conversion rates correspond to conversion tract initiation rates raging from 1 to 50 times the per-base pair crossover rate taken from `stdpopsim`.

Our baseline per-base pair gene conversion rates are equal to 6.55 *×* 10*^−^*^6^, 8.98 *×* 10*^−^*^6^, and 4.03 *×* 10*^−^*^7^ in humans, *D. melanogaster*, and *A. thaliana*, respectively. The value for humans very closely matches a recent estimate (5.9 *×* 10*^−^*^6^ from Williams *et al*. (2015)), as does the value for *D. melanogaster* (9.26 *×* 10*^−^*^6^ is the rate one obtains using the initiation and tract length estimates from Miller *et al*. (2016)). The value for *A. thaliana* is quite low compared to empirical estimates from F2 crosses (on the order of 5 *×* 10*^−^*^6^ Wijnker *et al*. (2013)). The reason for this discrepancy is that the *c* value obtained from `stdpopsim` accounts for selfing (following Huber *et al*. (2014) which assumed a 97% selfing rate). A value of *g* = 50 may therefore better correspond to the total gene conversion rate in *A. thaliana*, but if selfing is indeed quite prevalent, our value of *g* = 1 may better model the *effective* rate of gene conversion events. Thus, we can generally think of the baseline *g* = 1 simulations as indicative of the average effect of gene conversion on selective sweeps, and the higher values may only be relevant to loci/species where gene conversion occurs at especially high rates.

The beneficial allele’s selection coefficient, *s*, was set to 0.001, 0.01, or 0.1, and its dominance coefficient, *h*, was set to 0, 0.5, or 1.0. The time of onset of selection, *t_s_*, expressed in units of 4*N_A_* generations before the present (where *N_A_* is the ancestral population size of the simulated model), was set to 0.01, 0.05, 0.1, 0.5, or 1. We simulated 1000 replicates of all combinations of *g*, *s*, *h*, and *t_s_* for each demographic model examined (see below). If at any point during the simulation the beneficial mutation was lost, the simulation was reset to the point at which the mutation originally occurred.

We implemented this two-locus model using `SLiM` version 4.0.1, and simulating a chromosomal stretch where gene conversion tracts were allowed to initiate at the selected (*B*) locus only, such that the tracts would be extremely unlikely to extend to the linked (*A*) locus; this was done by setting the mean tract length to 2 and separating the two loci by a large distance (i.e. on the order of 10 kb or more). This strategy essentially ensured that any observed *aB* haplotypes were the result of gene conversion copying the *B* allele onto a *a* background.

### 2.2 Demographic models simulated via stdpopsim

For each species, we simulated sweeps under multiple demographic histories. This was done by using `stdpopsim` to generate a `SLiM` script for the desired demographic model, and then programmatically modifying that `SLiM` script to include a selective sweep at the desired time. For humans, we used three models: a model with a constant population size of *N* = 10, 000 individuals (`stdpopsim`’s default), the EUR population from Tennessen et al.’s (2012) two-population model as implemented by `stdpopsim` (`OutOfAfrica 2T12` in the `stdpopsim` catalog), and the AFR population from the same Tennessen et al. model. For *D. melanogaster*, we used two models: a constant-size model with *N* = 1, 720, 600 (`stdpopsim`’s default, from Li and Stephan (2006)), and the `African3Epoch 1S16` model, which is a one-population piecewise-constant 3-epoch model estimated by Sheehan and Song (2016). For *A. thaliana* the two models were a constant-size model with *N* = 10, 000 individuals (again, `stdpopsim`’ default), and `African3Epoch 1H18`, the 3-epoch one-population model from Huber et al. (2018). To accelerate the simulations, we divided population sizes and the number of generations in each epoch by a scaling factor, *Q*, and multiplied selection coefficients and gene conversion rates by this same factor. For humans and *A. thaliana* we set *Q* to 10, and for *D. melanogaster* we set *Q* to 100. We note that for large values of *s* it is possible that this rescaling effected our results (Uricchio and Hernandez, 2014), and thus we caution that, especially for the largest value of *s* examined here (0.1), our precise estimates of the fraction of sweeps that are pseudo-soft may be somewhat biased.

### 2.3 Simulation outcomes measured

For each replicate simulated under the models and parameter combinations described above, the simulation ended either when the beneficial allele reached fixation, or when the simulation reached the end of the specified demographic model (i.e. *t_s_ ×*4*N_A_* generations after the beginning of the sweep). At the end of the simulation, we asked whether the sweeping mutation had fixed, and if so, recorded the following outcomes: 1) whether or not the sweep was pseudo-soft at the population level (i.e. whether the *a* allele was present in the population), and if so 2) the population frequency of the *a* allele. We also took a random sample of 200 chromosomes from the population, and additionally asked: 3) whether or not the sweep was pseudo-soft in a random sample of 200 chromosomes taken from the population, and if so 4) the sample frequency of the *a* allele.

If the sweeping mutation had not fixed by the end of the simulation, we recorded the following outcomes: 1) the frequency of the the *B* allele at the end of the simulation, 2) whether or not the sweep was pseudo-soft in a random sample of 200 chromosomes (i.e. the *aB* haplotype was present in the sample), and if so, 3) the sample frequency of the *a* allele among those chromosomes with the *B* allele? (i.e. how soft was this partial sweep in the sample)? Note that because all simulation outcomes recorded depend only on the trajectory of the beneficial allele and the gene conversion events it experienced during its sojourn, the population’s evolutionary history prior to the sweep was not relevant, and therefore no burn-in period was required for these simulations. All code required for running these simulations, parsing their output, and visualizing results are available at https://github.com/SchriderLab/geneConvSweeps.

## 3 Results

### 3.1 Overview of two-locus simulation model and outcomes

We used forward-in-time simulations of positive selection acting on a *de novo* mutation to assess the impact that gene conversion is likely to exert on selective sweeps in natural populations. We examined this by performing. Importantly, these simulations did not include crossover events, but did allow gene conversion at the selected locus (Methods). We asked how often these sweeps were “softened” by conversion by recording how often recombinant haplotypes bearing the selected allele were present at the end of the simulation. Intuitively, the probability of observing such pseudo-soft sweeps depends on a number of parameters, including the total gene conversion rate per base pair (equal to the tract initiation rate multiplied by the mean tract length), the advantageous allele’s selection coefficient, *s*, and the dominance coefficient, *h*. We therefore simulated sweeps under a number of different combinations of values of each of these parameters.

The probability that a gene conversion event moves the beneficial allele onto one or more new genetic backgrounds that persist after the sweep may depend not only on the parameters described above, but also the demographic history of the population. For example, if a sweeping allele is present on multiple distinct haplotypes, the probability that all but one of these haplotypes are lost, resulting in a hard sweep, is elevated when the population size is reduced (Wilson *et al*., 2014). Relatedly, the history of changes to the population size (*N*) during selection, and thus the strength of selection *Ns*, will influence the sojourn of the adaptive allele. For both of these reasons, in populations whose size varies over time the outcome of a sweep in our model will depend on when the sweep began (*t_s_*). The histories of demographic changes vary considerably across species, as do gene conversion rates. Thus, to ensure that our simulated scenarios encompassed parameter ranges that model the dynamics of a variety of natural populations, we conducted our simulations using population size histories and gene conversion rates relevant to three species with very different sizes, demographic histories, and effective rates of recombination: humans, *D. melanogaster*, and *A. thaliana* (Methods). We simulated sweeps under multiple demographic histories for each species, and various values of *t_s_* as well (Methods). We report the results for simulations modeled after each species in turn below.

### 3.2 Gene conversion softens a minority of simulated sweeps in humans

We simulated three demographic models in humans: a constant-size population, a population recent experiencing exponential growth (the “African” subpopulation from the model presented in Tennessen *et al*. (2012), hereafter referred to as the AFR model), and a population experiencing two contractions followed by exponential growth and a more recent phase of accelerated growth (the “European” model from Tennessen *et al*. (2012), or EUR). We consider sweeps to be pseudo-soft if the beneficial allele is found on multiple haplotypes in a sample of 200 individuals drawn from the population at the conclusion of the simulation (although similar results are obtained if we examine the entire population; see Table S1). We summarize the simulation results for the constant-size model in Supplementary Figure 1. In this scenario, we observe that for a relative gene conversion rate of *g* = 1, which roughly corresponds to the rate in humans (Methods), between 0 and 17% of sweeps are pseudo-soft, depending on the timing of the sweep (*t_s_*), and the selection and dominance coefficients (*s*, and *h*, respectively). The role of *t_s_* in this model stems from the fact that the most recent sweeps have not had adequate time to reach high frequencies, and therefore have had fewer opportunities to be softened by gene conversion. Thus, let us first consider the oldest sweeps (*t_s_* = 1 *×* 4*N_A_* generations ago, where *N_A_* is the population size in the oldest epoch of the demographic model under consideration; bottom row of Figure 2), as the vast majority of sweeps (*>* 97%) at this time had reached fixation prior to sampling for all parameter combinations examined. Here, we find that sweeps with smaller selection coefficients were most likely to be softened: 14–17% of sweeps, depending on the dominance coefficient, were pseudo-soft when *s* = 0.001, but only 7–12% were pseudo-soft when *s* = 0.01, and 3–7% were pseudo-soft when *s* = 0.1. For each value of *s*, sweeps with lower dominance coefficients were slightly more likely to be pseudo-soft. Our observation that sweeps with lower *s* and *h* are softer is intuitive in that we expect sweeps that take longer to reach fixation to will experience more gene conversion events. We observed qualitatively similar results when *t_s_* = 0.5.

**Figure 2:**
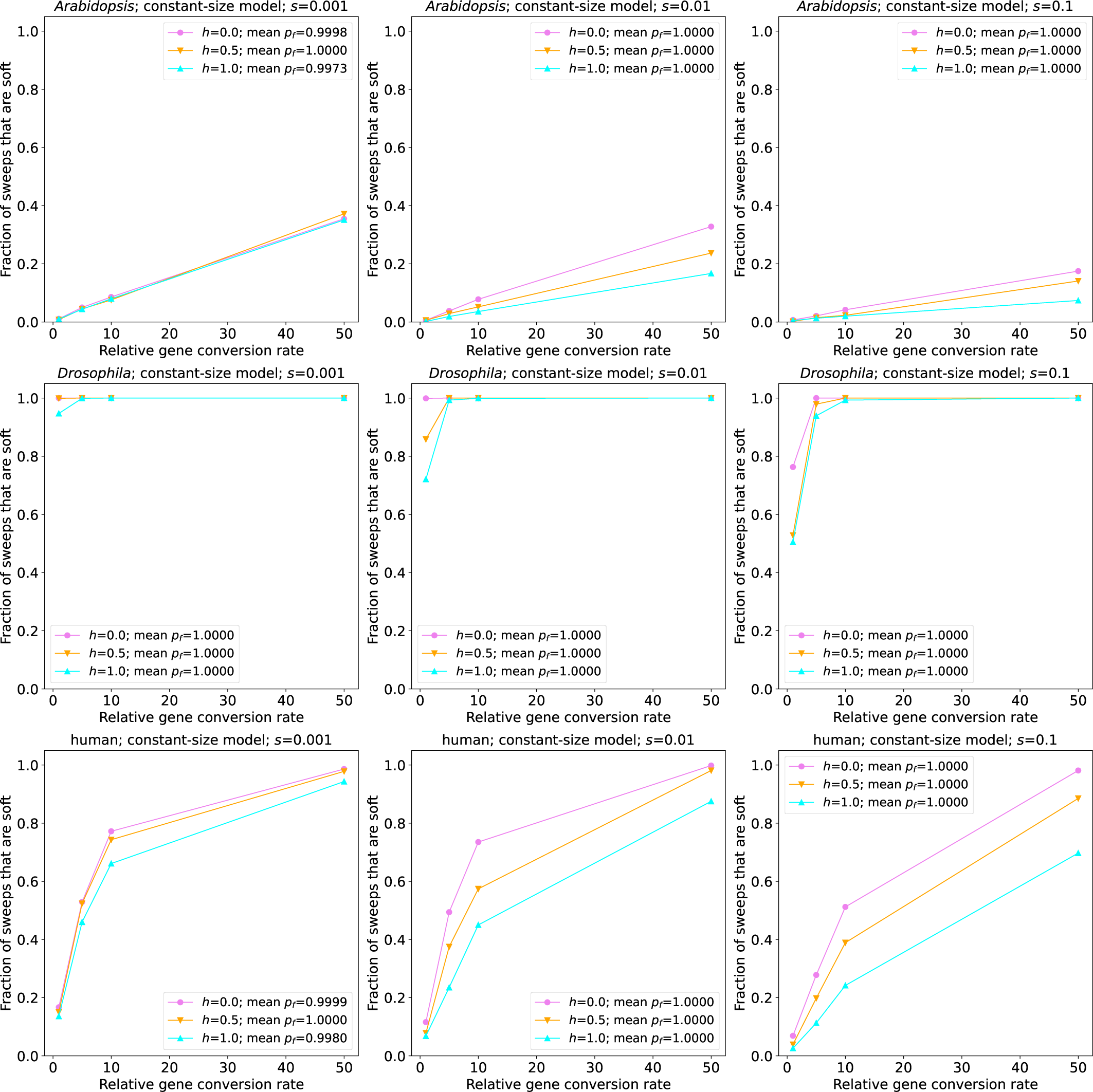
The fraction of sweeps that are pseudo-soft because of gene conversion in simulations of the constant-size *A. thaliana*(top row), *D. melanogaster* (middle row), and human populations (bottom row). Each panel shows the fraction of pseudo-soft sweeps at the time of sampling (*n* = 200 chromosomes) for a given combination of the selection coefficient (*s*) and simulated species, with the results for different dominance coefficients (*h*) shown in different colors as specified in the insets. For all simulations shown in this figure *t_s_* was set to 1.0, meaning that the simulation ran for up to 1.0 *×* 4*N* generations ago, at which point a sample of *n* = 200 chromosomes was taken from the population. If the sweep fixed prior to this sampling time, as was the case for the vast majority of replicates represented in this figure, the sample was instead taken at the time of fixation. The insets also show the average final frequency of the advantageous allele (*p_f_*) for each value of *h*.

When we examine sweeps that began far more recently, (i.e. *t_s_ ≤* 0.05), we instead observe that additive and dominant sweeps are more likely to be pseudo-soft when *s ≤* 0.01. For example, when *t_s_* = 0.05 and *s* = 0.001, 4.8% of sweeps are pseudo-soft when *h* = 1 versus 3.7% when *h* = 0. When *s* increases to 0.01, these fraction of pseudo-soft sweeps for *h* = 1 increases to 10.3%, while the fraction for *h* = 0 remains very low (3.7%). These more counter-intuitive results are simply a result of the much larger fraction of sweeps with low *s* and *h* that are still at low frequency at the time of sampling when *t_s_* is low. For instance, for the *t_s_* = 0.05 and *s* = 0.01 case described just above, the sweeping allele had reached a frequency of 0.91 on average in the *h* = 1 simulations, but only 0.16 when *h* = 0. Across all combinations of *s*, *h*, and *t_s_*, we observe that as the relative gene conversion rate increases, the fraction of sweeps that are pseudo-soft becomes far more appreciable, with an increase in *g* from 1 to 5 resulting in a 4.5-fold increase in the fraction of pseudo-soft sweeps, and the majority of sweeps being pseudo-soft when *g* = 50 for all cases where *t_s_ ≥* 0.01.

Next, we simulated sweeps occurring in two non-equilibrium demographic histories previously estimated from human population genetic data (Tennessen et al.’s AFR and EUR models noted above). In these scenarios, because the population size, *N*, changes over time, *t_s_* now exerts an effect beyond whether or not the sweeping allele has reached fixation prior to sampling: sweeps occurring at times where *N* is small will have lower values of *Ns*, a composite parameter indicating the strength of selection relative to drift, than sweeps occuring when *N* is larger. In Figure 3 and Supplementary Figure 2, we show results from simulations where we sampled individuals from the AFR and EUR populations, respectively. These results are similar to those from our constant-size model, with again between 0 and slightly under 20% of sweeps being softened when *g* = 1. The one notable difference was that when *t_s_* = 0.1, the EUR model yields a lower fraction of pseudo-soft sweeps than the constant-size and AFR models when *s* = 0.001 or 0.01. This is expected because a population contraction occurred during the sojourn of most of these sweeps, causing a loss of variation potentially removing recombinant haplotypes that might otherwise have persisted throughout the sweep (Wilson *et al*., 2014). As in our constant-size model, when we increase the gene conversion rate for sweeps under the AFR and EUR models we observe increasingly large fractions of pseudo-soft sweeps.

**Figure 3:**
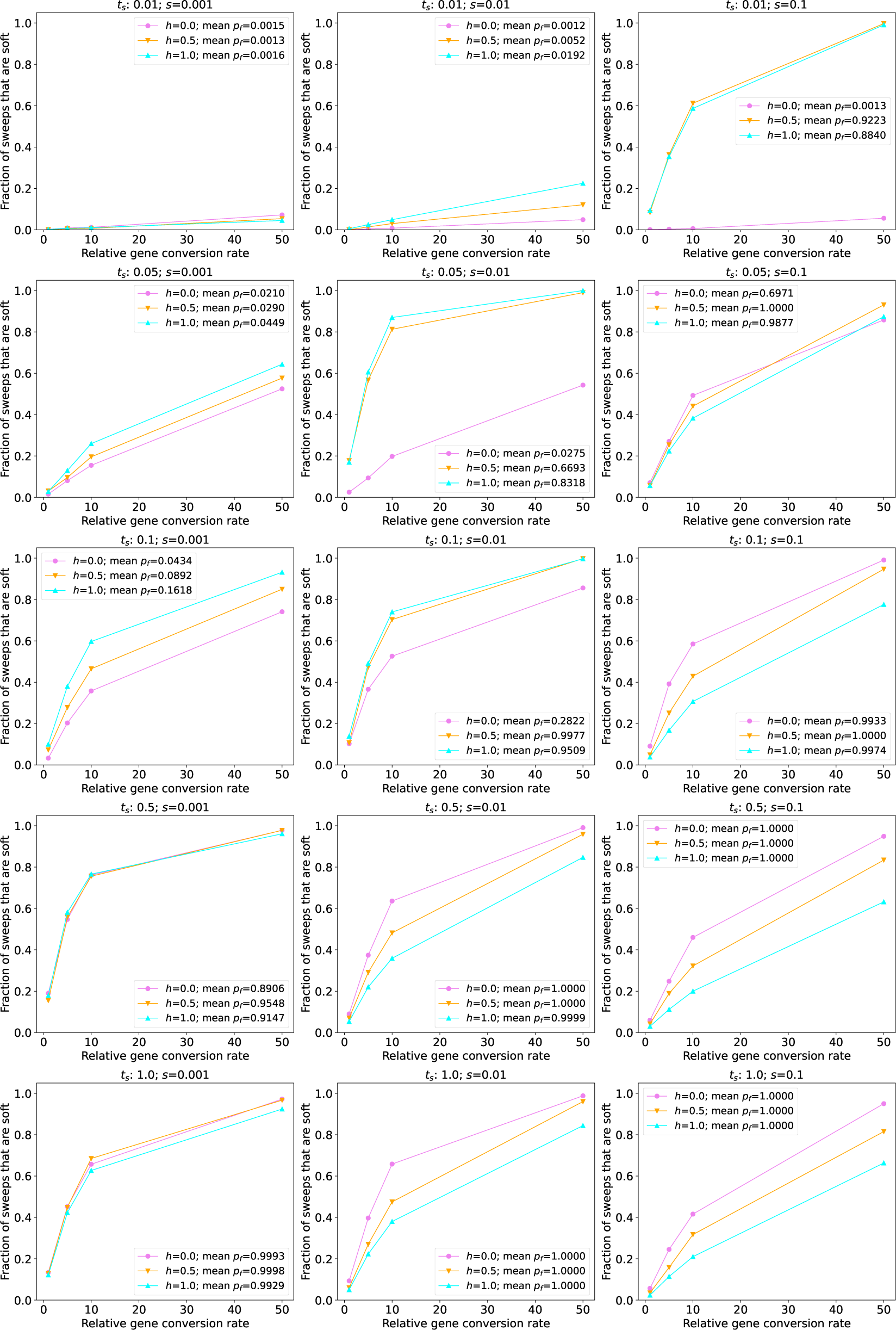
The fraction of sweeps that are pseudo-soft for simulations under the human AFR model (Tennessen *et al*., 2012) The fraction of pseudo-soft sweeps at the time of sampling (*n* = 200 chromosomes) for a given combination of the selection coefficient (*s*) and time since the start of the sweep (*t_s_*) are shown in the inset of the appropriate panel, with the results for different dominance coefficients (*h*) shown in different colors as specified in the insets. Because for more recent sweeps there was often not sufficient time for the sweeping allele to reach fixation, the insets also show the average final frequency of the advantageous allele (*p*_f_).

### 3.3 Gene conversion softens the majority of sweeps in simulated *D. melanogaster* populations

We next assessed the impact of gene conversion on sweeps in a larger population. Specifically, we simulated two scenarios modeled after *D. melanogaster* populations (see Methods for details): one with a constant population size, and the other following the demographic model from Sheehan and Song (2016), which contains a protracted population bottleneck. When simulating sweeps under the constant-size model, we observe trends that are similar to those seen for the human models (see the middle row of Figure 2 for sweeps with *t_s_* = 1, all of which reached fixation, and Supplementary Figure 3 for all values of *t_s_*). First, the fraction of sweeps that are pseudo-soft increases as selection gets weaker. Second, as the dominance coefficient decreases sweeps are more likely to be pseudo-soft, except when the sweep has not yet reached high frequency as is the case when *s* = 0.001 and *t_s_* = 0.01. However, the magnitude of gene conversion’s impact is far greater in the simulated *D. melanogaster* populations than observed in the human models: for most combinations of *t_s_*, *h*, and *s*, the majority of sweeps are pseudo-soft. Indeed, when *s* = 0.001 and *t_s_* = 0.05, all simulated sweeps were softened by gene conversion when *h* = 0.5 or 0.0, and *∼* 95% were softened when *h* = 1.0. For higher selection coefficients the majority of sweeps were pseudo-soft as well, although this fraction did vary according to the dominance coefficient. Even in the scenario where there is the least amount of time for gene conversion events to soften a sweep, i.e. sweeps acting on a dominant and strongly beneficial mutation (*h* = 1.0 and *s* = 0.1), roughly half of all sweeps were pseudo-soft (although this estimate may have been affected by our parameter rescaling as described in the Methods).

Next, we examined the effect of gene conversion on sweeps occurring throughout a 3-epoch bottleneck model estimated by Sheehan and Song (2016). For the most recent and the oldest sweep times simulated, our results under this model are very similar to those under the constant-size model (Figure 4). However, because this model experiences a bottleneck between 0.077 *×* 4*N_A_* and 0.84 *×* 4*N_A_* generations ago, for *t_s_* = 0.1 and *t_s_* = 0.5 the sweep occurs in the bottlenecked population and thus the softening effect of gene conversion is diminished. For both this model and the constant-size model, we again see that as the gene conversion rate increases, the fraction of pseudo-soft sweeps grows and eventually reaches 1.0. We expect that the majority of the *D. melanogaster* genome will not have such high rates of gene conversion, but even for *g* = 1 (which roughly corresponds to the average rate in *D. melanogaster*), for most combinations of *s*, *h*, and *t_s_* examined here, the majority of sweeps in *D. melanogaster* are softened by gene conversion.

**Figure 4:**
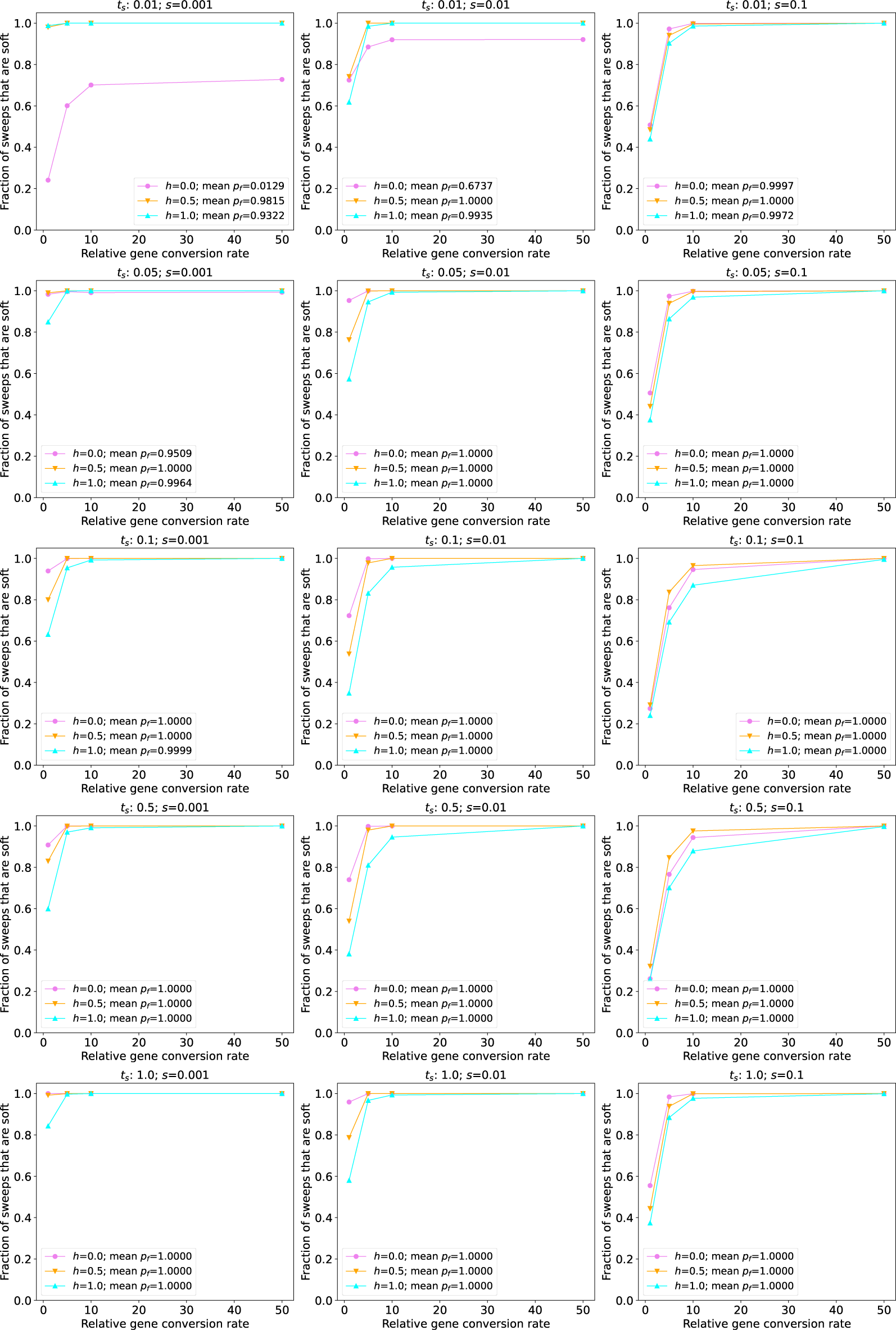
The fraction of sweeps that are pseudo-soft for simulations under the *Drosophila* 3-epoch model from (Sheehan and Song, 2016). The fraction of pseudo-soft sweeps at the time of sampling (*n* = 200 chromosomes) for a given combination of *s* and the time since the start of the sweep (*t_s_*) are shown in the appropriate panel, with the values for different dominance coefficients (*h*) shown as different colors as specified in the inset. The average final frequency of the advantageous allele (*p_f_*) for a given parameter combination is also shown in the inset. Note that the vast majority of sweeps were at or near fixation by the time of sampling except for the most recent sweeps examined (*t_s_* = 0.01). For *t_s_* = 0.01, recessive sweeping alleles with *s* = 0.001 were generally still at very low frequency (*p_f_* = 0.01 on average) at the time of sampling.

### 3.4 Gene conversion has a minimal impact on sweeps in simulated *A. thaliana* populations

Finally, we investigated the role of gene conversion in simulated populations of *A. thaliana*, a species that undergoes a high rate of self-fertilization (Platt *et al*., 2010), and therefore should have fewer opportunities for gene conversion during a sweep. We began by simulating a constant-size population of 10,000 individuals, `stdpopsim`’s default size for this species. As shown in the top row of Figure 2 (for sweeps with *t_s_* = 1, nearly all of which reached fixation) and Supplementary Figure 4 (for all values of *t_s_*), a very small fraction of sweeps are pseudo-soft in this model, and increasing the gene conversion rate has a smaller effect than in the human simulations. This may be expected given the relatively small effective population combined with the very low rate of effective gene conversion used to account for selfing (Methods). We therefore examined the 3-epoch South Middle Atlas model from Huber et al. (2018), which in spite of a bottleneck contains substantially larger effective population sizes throughout: 161,744 in the first epoch, 24,076 in the second epoch, and 203,077 in the final epoch. Under this model we observe a larger fraction of soft sweeps, although the highest observed for our baseline gene conversion rate is 11.5%. Thus, even a species that undergoes frequent selfing may experience pseudo-soft sweeps via gene conversion, although the fractions observed in these simulations are far below those in the *D. melanogaster* simulations. We also found that under the South Middle Atlas model, as the gene conversion rate increases, the fraction of pseudo-soft sweeps increases at a greater rate than under the constant-size model.

### 3.5 The frequency of recombinant haplotypes in pseudo-soft sweeps

As described above, our simulations show that allelic gene conversion has the potential to produce sweeps where the beneficial mutation is found on multiple distinct haplotypes. This suggests that pseudo-soft sweeps caused by gene conversion, much like true soft sweeps, may provide the means for a larger fraction of diversity present prior to selection to escape the sweep than expected under a hard sweep. To more directly address this possibility, we examined our simulated pseudo-soft sweeps and asked how many recombinant haplotypes were present upon fixation of the sweeping allele. Specifically, for each simulation replicate in which a sweep reached fixation and was categorized as pseudo-soft, we obtained the fraction of chromosomes that exhibited the *aB* haplotype.

We found that, conditioning on a pseudo-soft sweep having occurred, a substantial fraction of haplotypes in a sample taken immediately post-fixation were recombinants. For example, in our simulations under the *D. melanogaster* constant-size model, the median recombinant haplotype frequency ranges from 0.095 when *s* = 0.001, 0.015 when *s* = 0.01, to 0.005 when *s* = 0.1 for sweeping additive mutations. We observe similar recombinant frequencies for dominant sweeps under this model. For recessive sweeps, we observe much higher frequencies of the recombinant haplotype: 0.62 when *s* = 0.001, 0.155 when *s* = 0.01, and 0.025 when *s* = 0.1. Under the human constant-size model, we observe slightly higher median recombinant fractions for the dominant and additive cases than in the *D. melanogaster* model: 0.26, 0.075, and 0.023 for dominant sweeps (with *s* = 0.001, *s* = 0.01, and *s* = 0.1, respectively), and 0.17, 0.035 and 0.02 for additive sweeps. However, for recessive sweeps we observe much lower recombinant frequencies in humans than in *D. melanogaster* (0.15, 0.05, and 0.025, respectively, at increasing values of *s*), except for *s* = 0.1 where we see no difference between the two species. For both species, the frequency of recombinant haplotypes increases as the gene conversion rate increases, eventually approaching 1 in some cases, meaning that the original, unconverted sweeping haplotype has been eliminated by the time the sweeping allele has reached fixation. Note that even though only the recombinant haplotype was present after the sweep, we still considered these cases to be pseudo-soft because these sweeps involved multiple independent conversion events that almost certainly would have resulted in the advantageous allele being present on multiple genetic backgrounds had our simulations incorporated ancestral variation segregating at multiple linked loci at the onset of selection.

Interestingly, the phenomenon of the unconverted haplotype being completely eliminated at the expense of the recombinant haplotype was not exclusively observed when the gene conversion rate was exceptionally large. For example, even at a relative gene conversion rate of 1, 30% of our simulated *D. melanogaster* sweeps with *h* = 0 and *s* = 0.001 resulted in the loss of the unconverted haplotype, although this fraction dropped to *∼*1% for dominant sweeps. Intriguingly, sweeps in the human model with *s* = 0.001 displayed the opposite trend, with higher median recombinant haplotype frequencies for dominant sweeps than recessive sweeps (Figure 6), and a slightly higher fraction of sweeps where only recombinant haplotypes survived the sweep when *h* = 1 (16% of sweeps had only the *aB* haplotype) than when *h* = 0 (11% of sweeps).

**Figure 5:**
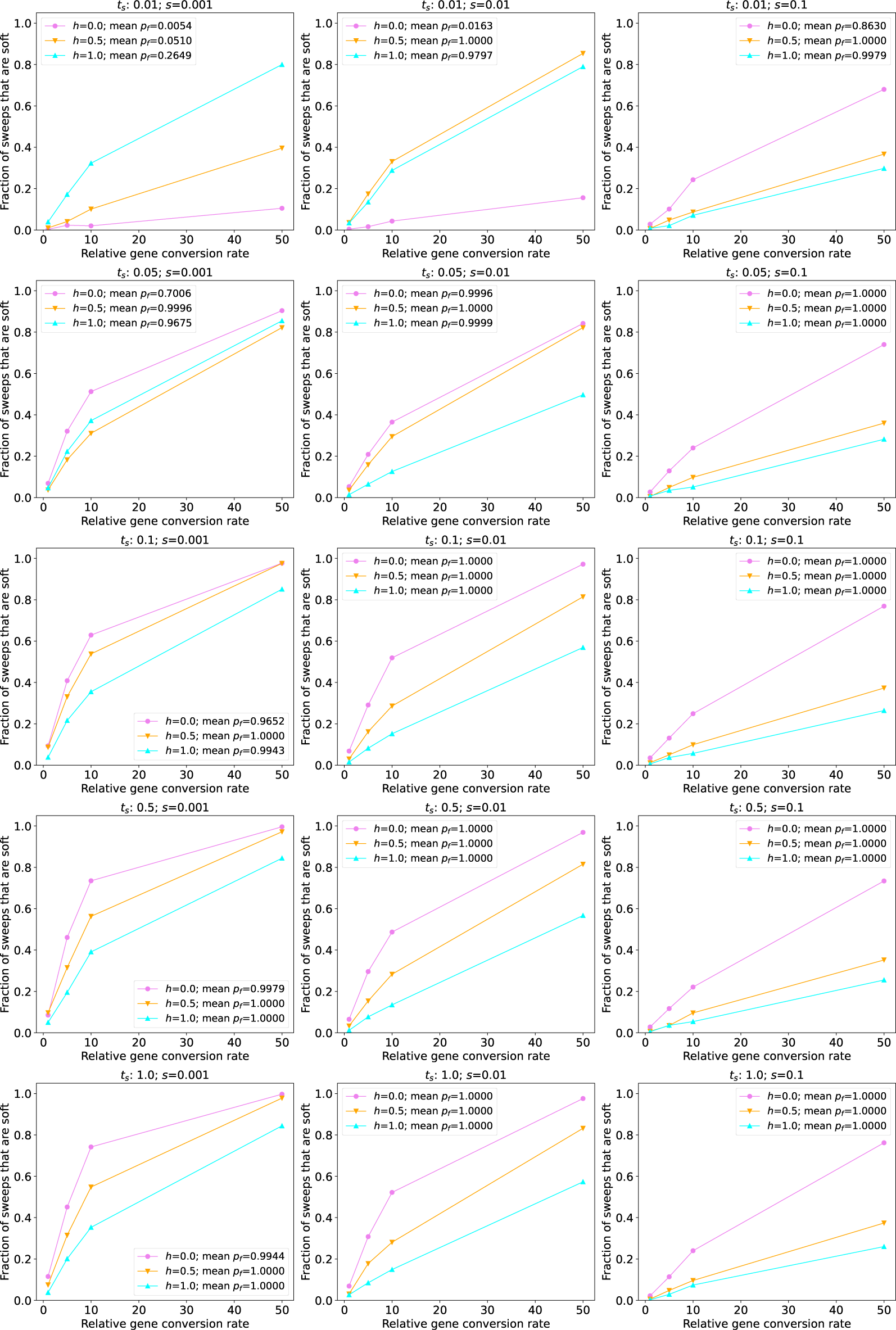
The fraction of sweeps that are pseudo-soft for simulations under the *Arabidopsis* 3-epoch South Middle Atlas model (Huber *et al*., 2018). The fraction of pseudo-soft sweeps at the time of sampling (*n* = 200 chromosomes) for a given combination of *s* and time since the start of the sweep (*t_s_*) are shown in the appropriate panel, with the values for different dominance coefficients (*h*) shown as different colors as specified in the inset. The average final frequency of the advantageous allele (*p_f_*) for a given parameter combination is also shown in the inset. Note that the vast majority of sweeps were at or near fixation by the time of sampling except for the very recent sweeps examined (beginning at *t_s_* = 0.01 or *t_s_* = 0.05). For *t_s_* = 0.01, recessive sweeping alleles with *s* = 0.001 and *s* = 0.01 were generally still at very low frequency (*p_f_* = 0.01 and *p_f_* = 0.02 on average, respectively) at the time of sampling.

**Figure 6:**
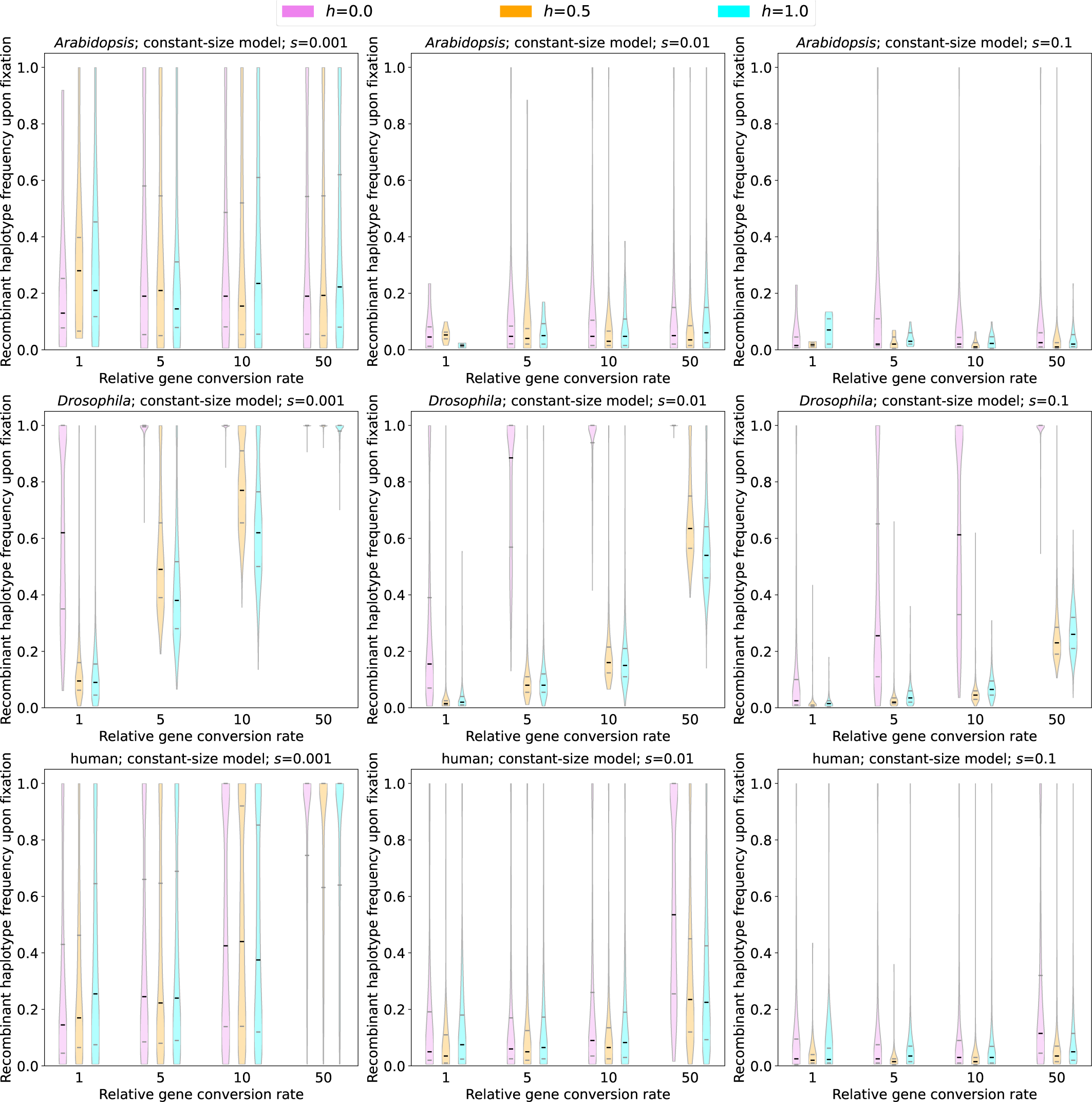
The sample frequency of recombinant (*aB*) haplotypes upon fixation for sweeps under each species’ constant-size model. Violin plots show the distribution of recombinant haplotype frequencies, illustrating the degree of softness produced by gene conversion under a given parameter combination, with the violin color corresponding to the dominance coefficient as specified in the inset. Each row shows results for a different species (*A. thaliana*, *D. melanogaster*, and human, going from top to bottom), and each column shows results for a different selection coefficient (*s* = 0.001, *s* = 0.01, and *s* = 0.1 going from left to right). All results were taken from simulations with *t_s_* = 1.

We note that our results in our *A. thaliana* constant-size model are generally comparable to those in our human simulations with a relative gene conversion rate of 1, but as the gene conversion rate increases the *A. thaliana* recombinant frequencies do not increase at the same rate as in humans. Although we caution that the estimates in the *A. thaliana* models are based on a much smaller number of pseudo-soft sweeps than for other species, we again found that the recombinant frequency upon fixation increased as the gene conversion rate increased and as *s* decreased.

Next, we examined the immediate post-fixation frequency of recombinant haplotypes created by gene conversion under each of the simulated non-equilibrium demographic histories. Unlike the constant-size histories considered above, in these models the softness of a completed sweep may depend on the precise timing of its sojourn vis-a-vis population size changes. We therefore show the recombinant frequencies for all fixation times in Supplementary Figures 5–8. Generally, we observe lower frequencies of recombinant haplotypes in these models, perhaps in part because of the effect of population bottlenecks, although in most cases the difference between the constant-size and non-equilibrium models for a given species were fairly small. One notable exception is the set of recessive sweeps with *s* = 0.001 in our *D. melanogaster* simulations: although in our constant-size simulations we observed median recombinant frequency of *>* 60%, across values of *t_s_* in our non-equilibrium simulations the median recombinant frequency never eclipsed 40% (Supplementary Figure 5). We also found that the non-equilibrium *A. thaliana* simulations yielded higher median recombinant frequencies than those observed under the constant-size model, while noting that the latter are based on a very small number of pseudo-soft sweeps, and a more appreciable number of pseudo-soft sweeps occurred under the non-equilibrium model (Figure 5).

## 4 Discussion

Searching for signatures of positive selection is a challenging but important task, as such studies can reveal loci responsible for recent adaptation (Sabeti *et al*., 2002; Nielsen *et al*., 2005; Pickrell *et al*., 2009; Garud *et al*., 2015). In recent years, there has been increasing interest in inferring not only the targets of positive selection, but also the mode of adaptation (Jones *et al*., 2012; Garud *et al*., 2015; Feder *et al*., 2016; Schrider and Kern, 2017; Xue *et al*., 2021)—does adaptation proceed through classic “hard” sweeps on *de novo* mutations or via “soft sweeps” on standing variation or recurrent mutations? The answer to this question has an important implication for the rate of adaptation in natural populations: if soft sweeps are common, it may imply that the population is not mutation limited, meaning that adaptation to a new selective pressure may proceed immediately rather than requiring a long waiting time for a beneficial substitution to occur (Orr and Betancourt, 2001; Hermisson and Pennings, 2005; Karasov *et al*., 2010).

The defining feature of soft sweeps is that after the sweep has completed, the haplotypes present upon fixation share a common ancestor that occurred before the onset of selection (Hermisson and Pennings, 2017). However, previous studies have shown that gene conversion can yield a similar outcome by copying the advantageous allele onto new haplotypes during a sweep of a single-origin *de novo* mutation, a phenomenon we refer to here as a “pseudo-soft sweep”. First, Jones and Wakeley (2008) showed that gene conversion events occurring during a sweep can produce signatures of LD that differ from those expected under a hard sweep. Further, Schrider et al. 2015 showed that classic sweeps with gene conversion may generate patterns of diversity that resemble those of soft sweeps. However, to date no studies have asked how often pseudo-soft sweeps may occur in more realistic models that contain demographic histories and gene conversion rates estimated from natural populations.

We simulated selective sweeps on single-origin *de novo* mutations using demographic models and gene conversion rates estimated from humans, *D. melanogaster*, and *A. thaliana*. We found that gene conversion may often produce pseudo-soft sweeps, especially for smaller values of *s* and *h* and for larger population sizes. In other words, parameters that increase the number of generations for the beneficial allele to fix result in more cumulative opportunity for gene conversion to copy the allele onto new genetic backgrounds. By the same token, demographic changes can counteract the softening effect of gene conversion. For example, the probability of a pseudo-soft sweep is generally lower under the human EUR model from Tennessen et al. (2012) then under the AFR and constant-size human models (compare Supplementary Figure 2 to Figure 3 and Supplementary Figure 1). This may be due in part to the hardening effect of population bottlenecks causing all but one haplotype with the adaptive allele to be lost (Wilson *et al*., 2014), and also a result of smaller population sizes having smaller absolute sojourn times and thus fewer chances for gene conversion.

Not only can gene conversion allow more than one haplotype to survive the sweep, the frequencies of recombinant haplotypes often account for a sizeable portion of the sample. This outcome also depends on the selective and demographic parameters of the population. For example, *h* appears to again play an important role, with much larger frequencies of the *aB* haplotype upon fixation for recessive sweeps than additive or dominant sweeps in our *D. melanogaster* simulations. Indeed, in many *D. melanogaster* simulations of recessive sweeps with realistic gene conversion rates the original haplotype bearing the advantageous allele is lost, and only haplotypes that have acquired the adaptive allele through gene conversion are present at the time of fixation. However, the effect of dominance can vary across models, as exemplified by our human simulations with *s* = 0.001, wherein the recombinant haplotype frequency upon completion of the sweep was higher when *h* = 1 than *h* = 0. In addition, we emphasize that the *aB* haplotype frequency can be fairly high (*∼*10% or greater) even for additive or dominant sweeps in all three species modeled here when *s* = 0.001. Thus, when realistic levels of gene conversion do produce pseudo-soft sweeps, the reduction in linked genetic diversity should be considerably less severe than that of hard sweeps. At higher rates of gene conversion, the degree to which sweeps are softened is dramatically increased in our human and *D. melanogaster* simulations, with the original sweeping haplotype lost in the majority of cases—we expect that this would be the norm only in genomic regions with exceptionally high rates of allelic gene conversion.

Perhaps our most striking finding is that, in simulations with population sizes and gene conversion rates similar to those estimated from *D. melanogaster*, the majority of sweeps appear to be soft despite the absence of recurrent mutation and/or selection on standing variation. We do note that, because we varied the timing of sweeps, we were able to observe the effect of the population bottleneck in Sheehan and Song’s (2016) model, finding that sweeps whose sojourn overlapped with this bottleneck had lower probabilities of being softened by gene conversion than those occurring either before or after the bottleneck (Figure 4). However, even for sweeps occurring during the bottleneck, the majority of were pseudo-soft except for those with *s* = 0.1 or dominant sweeps with *s* = 0.01. The main takeaway is that sweeps in large populations, especially those with smaller selection coefficients, should be softened by gene conversion. This is an intriguing result given that it has been argued that previous evidence for soft sweeps might instead be better explained by weak positive selection (Harris *et al*., 2018), which results in slower sweeps that leave more time for recombination to rescue flanking diversity, thereby lessening the impact of the hard sweep. Our finding appears to be at odds with this notion, at least for large populations with gene conversion rates similar to that of *D. melanogaster*, as slower sweeps due to weaker selection will be almost certainly softened by gene conversion in such populations. Any sweeps of strongly beneficial alleles that have occurred in *D. melanogaster*, however, are unlikely to have been hard sweeps, as the massive reduction in diversity produced under such a scenario (Garud *et al*., 2021) appears to be inconsistent with patterns of haplotypic diversity around the strongest selective sweep candidates in this species (Garud *et al*., 2015). Thus, neither hard sweeps via strong or weak selection appear to be a reasonable explanation for sweep signatures in *D. melanogaster*, and we conclude that sweeps that appear to be hard in this species should very much be the exception rather than the rule. On the other hand, our findings do make the biological interpretation of soft sweep signatures in this species, and those with similarly large populations, somewhat murkier, as we expect to observe apparent soft sweeps even if selection has acted on a single-origin *de novo* mutation. We also note that gene conversion could have an effect on sweeps in smaller populations as well, with pseudo-soft sweeps potentially being at most partially responsible to the lack of classic strong sweep signatures in humans (Hernandez *et al*., 2011), but we do not expect gene conversion’s role to be as prominent in humans as in species with larger populations.

Part of the reason for the controversy surrounding soft sweeps is that, because of their more subtle signature, detecting them with a low false positive rate is more difficult (Schrider and Kern, 2016). Thus, it would be convenient for sweep-detection efforts if positive selection rarely resulted in soft sweeps. However, we do not appear to be this fortunate—regardless of the mode of selection, signatures essentially indistinguishable from soft sweeps should be the norm in species with large population sizes and per-base pair gene conversion rates comparable to those measured in several animal species. It is also worth noting that our conclusions may not be limited to models with a single target of selection, as polygenic selection can in many cases produce selective sweeps (Thornton, 2019), which of course may also experience gene conversion. Thus, even if true soft sweeps never occur, although this appears unlikely as argued above, the presence of gene conversion necessitates methods capable of detecting the signatures of soft sweeps if we hope to be able to uncover loci responsible for recent adaptation. Continued advances on this front, such as machine learning methods that can detect soft sweeps with increasingly high accuracy (e.g. (Schrider and Kern, 2016; Kern and Schrider, 2018; Mughal and DeGiorgio, 2019; Lauterbur *et al*., 2023b; Arnab *et al*., 2023), are therefore essential if we wish to fully understand the importance of positive selection in recent evolutionary history and its role in shaping patterns of diversity.

## Acknowledgments

I thank Andrew Kern for feedback on the manuscript. This work was funded by the National Institutes of Health under award number R35GM138286.

## 5 Data Availability Statement

All code necessary to reproduce the work in this manuscript, including all simulations and downstream analysis, is available at https://github.com/SchriderLab/geneConvSweeps.

---

**Supplementary Figure 1:**
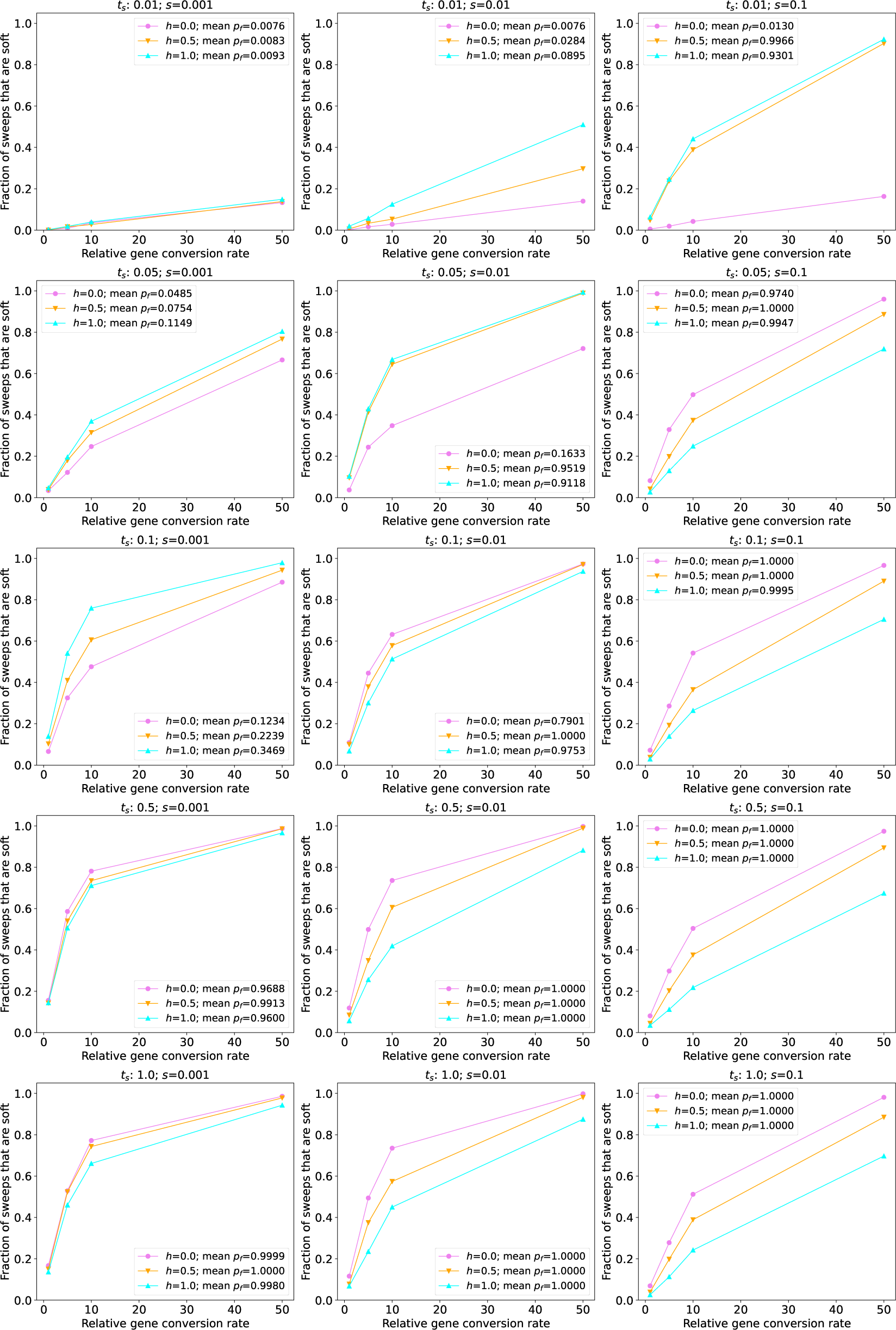
The fraction of sweeps that are pseudo-soft for the constant-population size human simulations. The fraction of pseudo-soft sweeps at the time of sampling (*n* = 200 chromosomes) for a given combination of the selection coefficient (*s*) and the time since the start of the sweep (*t_s_*) are shown in the appropriate panel, with the results for different dominance coefficients (*h*) shown as different colors as specified in the insets. Because for more recent sweeps there was often not sufficient time for the sweeping allele to reach fixation, the insets also show the average final frequency of the advantageous allele (*p_f_*).

**Supplementary Figure 2:**
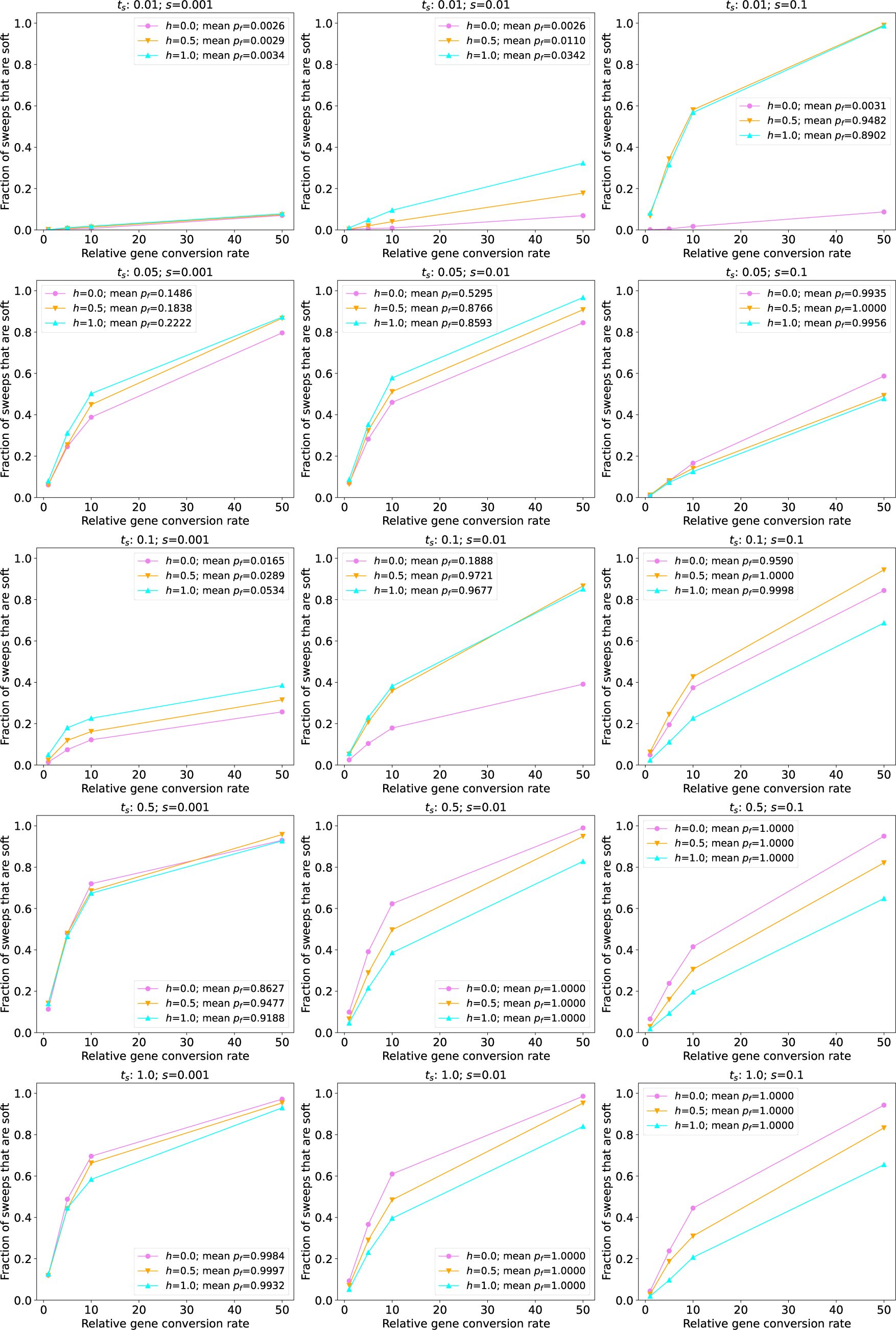
The fraction of sweeps that are pseudo-soft for simulations under the human EUR model. The fraction of pseudo-soft sweeps at the time of sampling (*n* = 200 chromosomes) under a given combination of *s* and the time since the beginning of the sweep (*t_s_*) are shown in the appropriate panel, with the results for different dominance coefficients (*h*) shown as different colors as specified in the inset. Because for more recent sweeps there was often not sufficient time for the sweeping allele to reach fixation, the insets also show the average final frequency of the advantageous allele (*p_f_*).

**Supplementary Figure 3:**
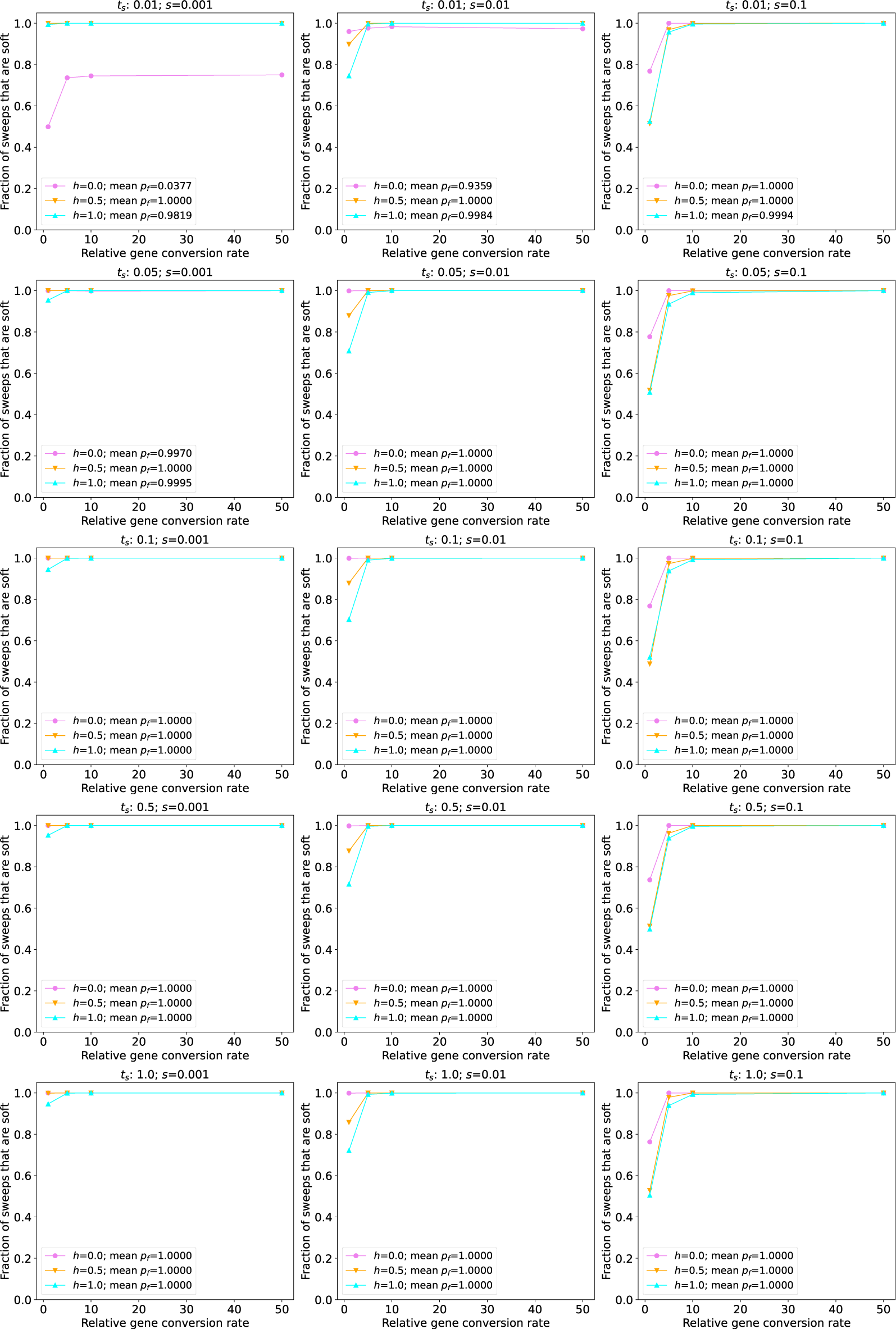
The fraction of sweeps that are pseudo-soft for the constant-population size *D. melanogaster* simulations. The fraction of pseudo-soft sweeps at the time of sampling (*n* = 200 chromosomes) under a given combination of *s* and the time since the beginning of the sweep (*t_s_*) are shown in the appropriate panel, with the results for different dominance coefficients (*h*) shown as different colors as specified in the insets. The average final frequency of the advantageous allele (*p_f_*) for a given parameter combination is also shown in the inset. Note that for the most recent sweeps examined (*t_s_* = 0.01), only 4% of recessive beneficial mutations with *s* = 0.001 had reached fixation at the time of sampling. However, for all other *t_s_* values examined, approximately all sweeps reached fixation prior to sampling, and therefore the results are roughly identical to one another across values of *t_s_* with the exception of the *s* = 0.001 and *h* = 0.0 case.

**Supplementary Figure 4:**
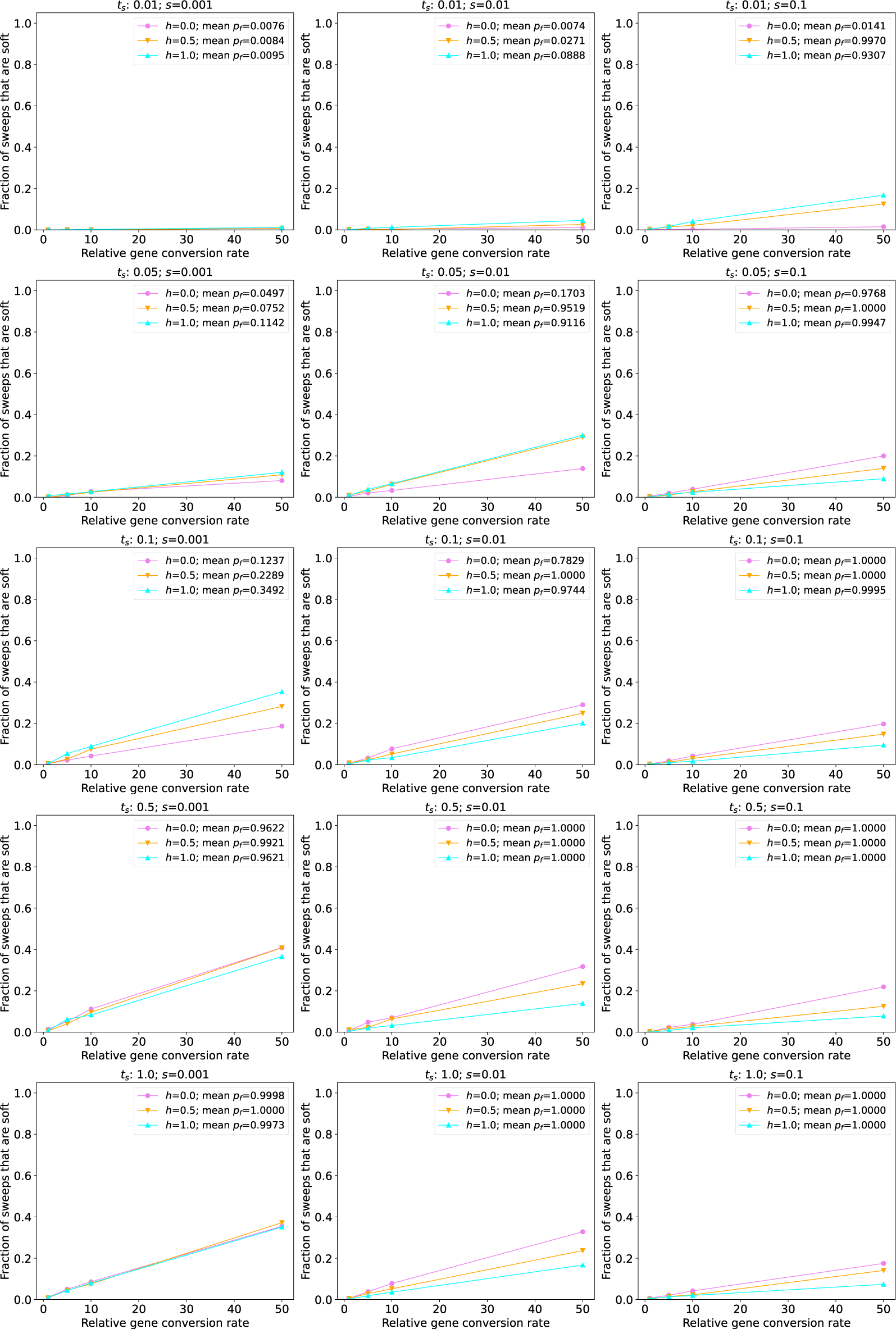
The fraction of sweeps that are pseudo-soft for simulations under the *Arabidopsis* constant-size model. The fraction of pseudo-soft sweeps at the time of sampling (*n* = 200 chromosomes) under a given combination of *s* and the time since the beginning of the sweep (*t_s_*) are shown in the appropriate panel, with the results for different dominance coefficients (*h*) shown as different colors as specified in the insets. Because for more recent sweeps there was often not sufficient time for the sweeping allele to reach fixation, the insets also show the average final frequency of the advantageous allele (*p_f_*).

**Supplementary Figure 5:**
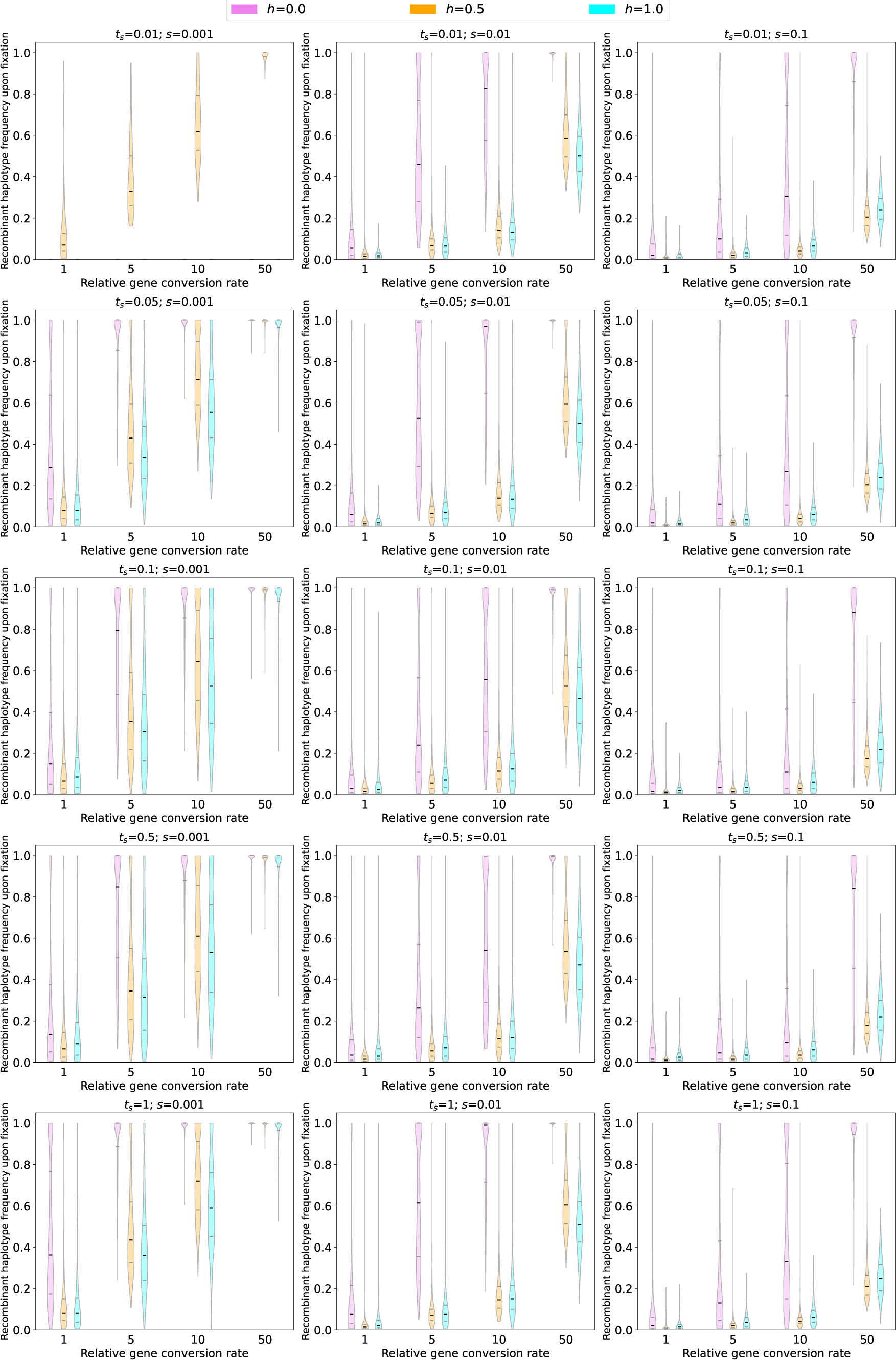
The sample frequency of recombinant (*aB*) haplotypes upon fixation for each combination of *s*, *h*, and *t_s_* under the *D. melanogaster* 3-epoch model (Sheehan and Song, 2016). Violin plots show the distribution of recombinant haplotype frequencies only for sweeps that had reached fixation at the time of sampling. For parameter combinations where no sweeps had fixed, no violins are shown.

**Supplementary Figure 6:**
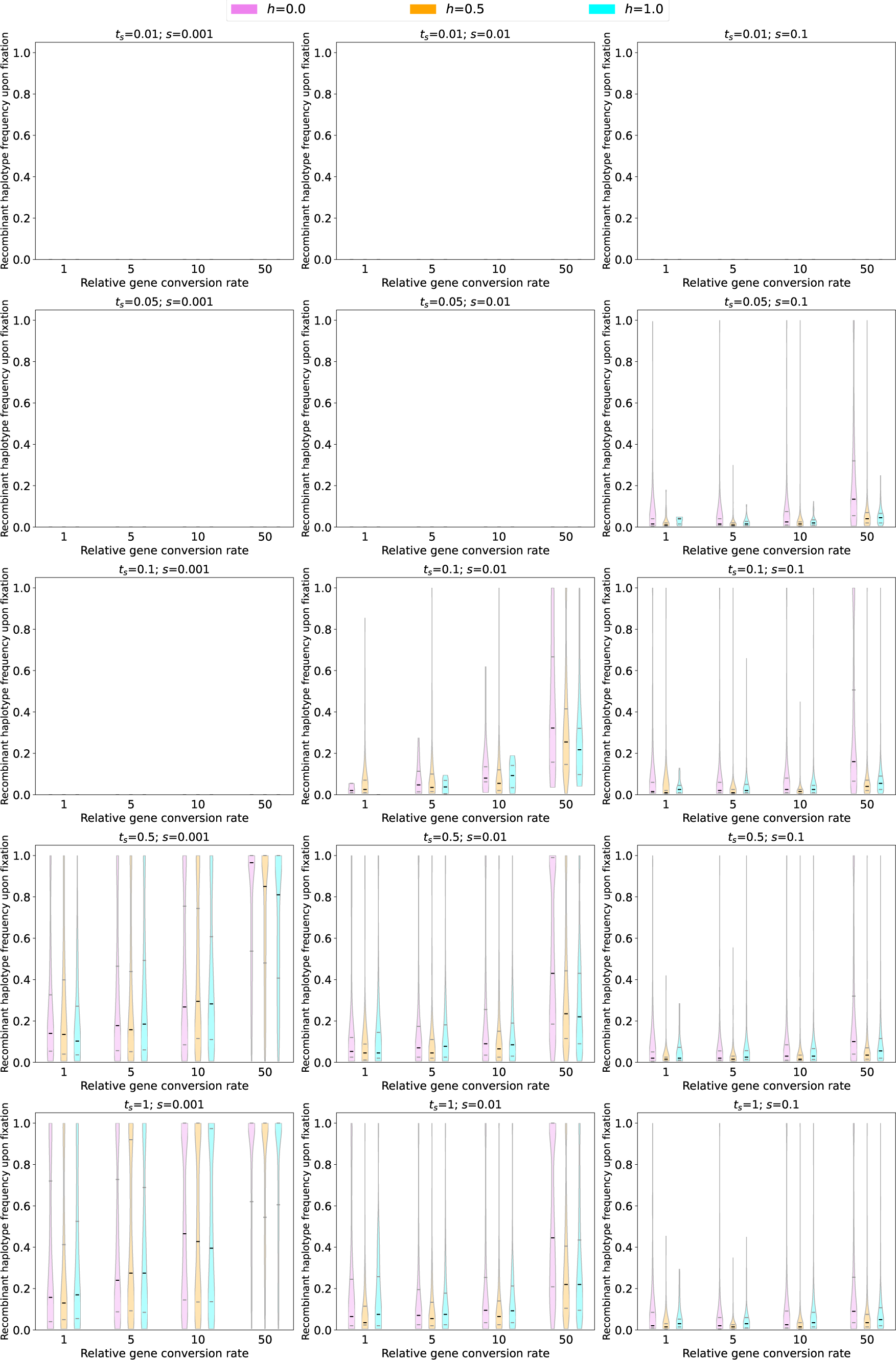
The sample frequency of recombinant (*aB*) haplotypes upon fixation for each combination of *s*, *h*, and *t_s_* under the human AFR model (Tennessen *et al*., 2012). Violin plots show the distribution of recombinant haplotype frequencies only for sweeps that had reached fixation at the time of sampling. For parameter combinations where no sweeps had fixed, no violins are shown.

**Supplementary Figure 7:**
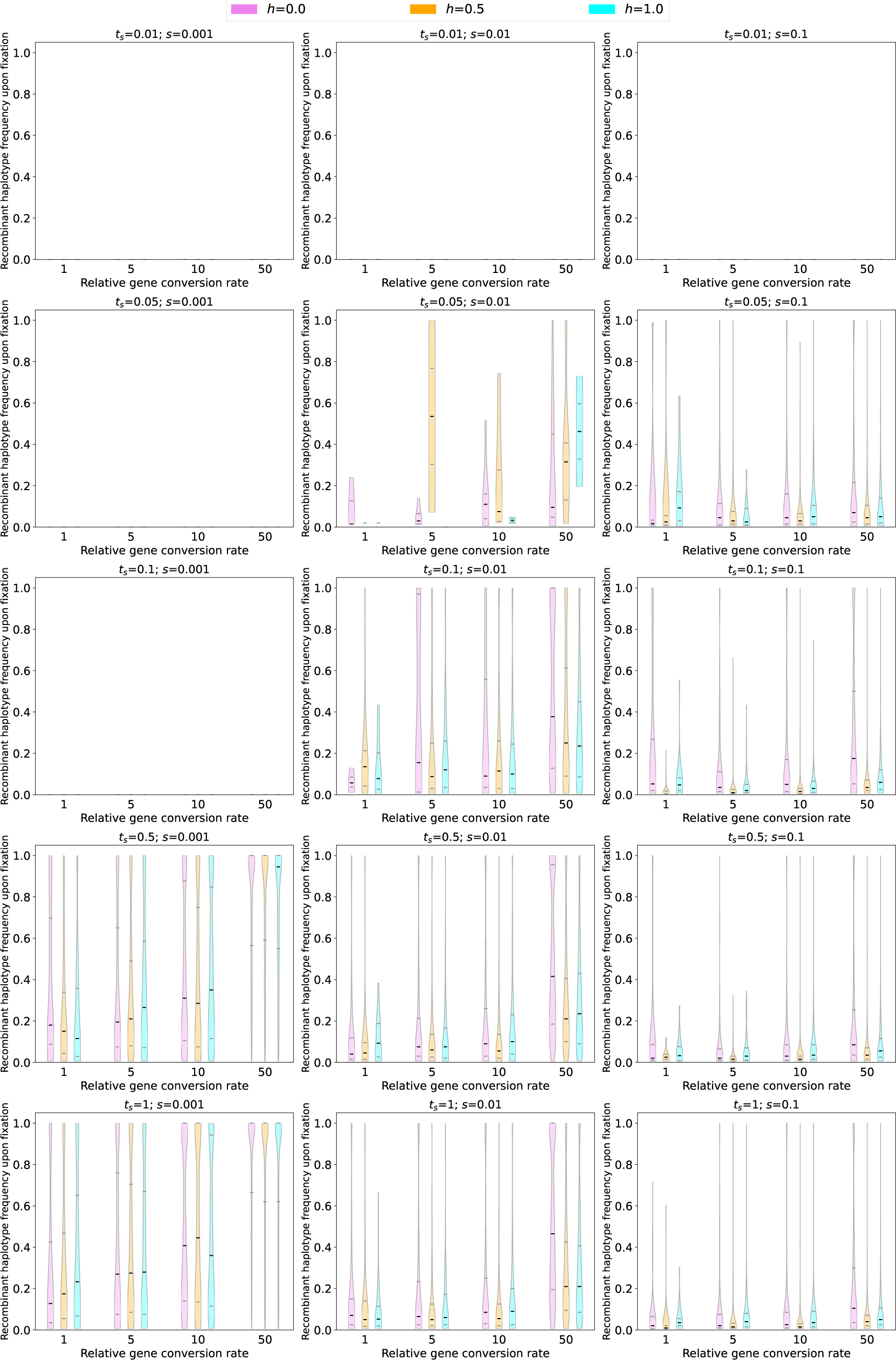
The sample frequency of recombinant (*aB*) haplotypes upon fixation for each combination of *s*, *h*, and *t_s_* under the human EUR model (Tennessen *et al*., 2012). Violin plots show the distribution of recombinant haplotype frequencies only for sweeps that had reached fixation at the time of sampling. For parameter combinations where no sweeps had fixed, no violins are shown.

**Supplementary Figure 8:**
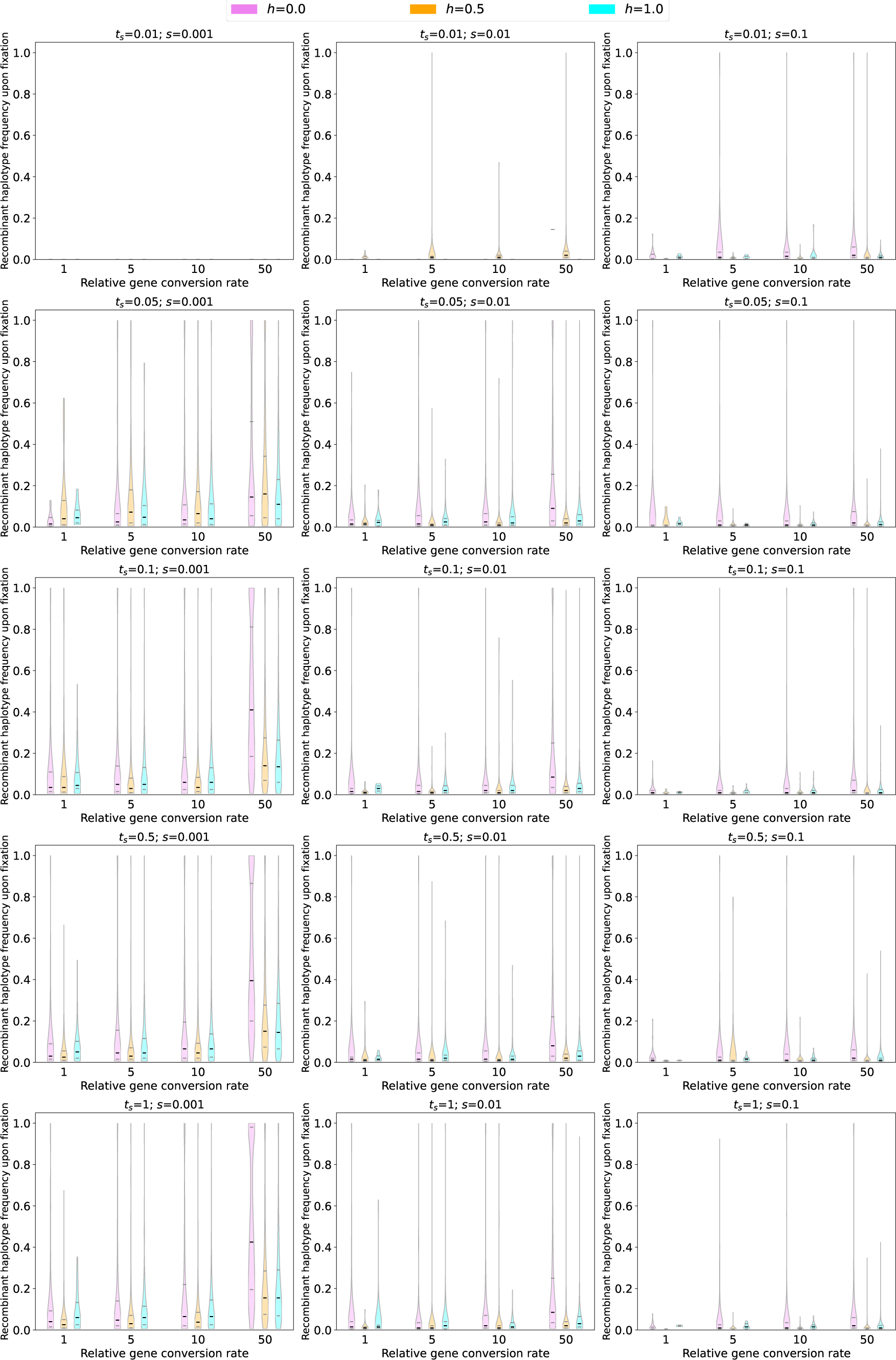
The sample frequency of recombinant (*aB*) haplotypes upon fixation for each combination of *s*, *h*, and *t_s_* under the *A. thaliana* 3-epoch model (Huber *et al*., 2018). Violin plots show the distribution of recombinant haplotype frequencies only for sweeps that had reached fixation at the time of sampling. For parameter combinations where no sweeps had fixed, no violins are shown.

